# Intraguild predation and life-history variation in a natural nematode predator-prey system

**DOI:** 10.64898/2026.04.02.716111

**Authors:** Desiree L. Goetting, Kamal Kaur Sarai, Penghieng Theam, Ralf J. Sommer, James W. Lightfoot

## Abstract

Predator-prey interactions are key drivers of behavioural and life-history evolution, as reciprocal selection can shape adaptations in both predators and prey. Yet, understanding how such interactions evolve in nature remains challenging because the ecological context in which they occur is often difficult to experimentally access. The nematode *Pristionchus pacificus* is a well-established model for studying predatory behaviour, but most experimental work has examined its predatory interactions with *Caenorhabditis elegans*, a species that *P. pacificus* is unlikely to encounter frequently in nature. Here, we investigate predation in a natural ecological context by focusing on nematode communities associated with beetles, the native necromenic habitat of *P. pacificus*. We identify *Oscheius myriophilus* as a species that co-occurs with *P. pacificus* in this niche and represents a natural prey species. Using established predatory assays, we show that *P. pacificus* actively attacks, kills, and consumes *O. myriophilus*. Strikingly, predation rates are significantly lower than those observed for *C. elegans*, consistent with the possibility that prolonged ecological association has favoured adaptations that reduce predation susceptibility. Supporting this interpretation, we identify distinct life-history traits in *O. myriophilus*, including a mixed reproductive strategy involving early oviposition followed by internal hatching of later progeny and matricide. Later developmental stages are more resistant to predatory attack, suggesting that retaining embryos within the maternal body may provide a protected environment in which offspring can develop while remaining buffered from predation. Together, our findings establish *P. pacificus* and *O. myriophilus* as a tractable system for investigating predator-prey interactions in a natural ecological context and provide a foundation for dissecting how predatory behaviour and prey life-history strategies evolve in response to one another.

## Introduction

Predator-prey interactions can be powerful drivers of evolution as they impose reciprocal selective pressures on interacting species. Predators may evolve traits that increase their ability to locate, capture, and consume prey, while prey experience selection for traits that reduce their susceptibility to attack. Such reciprocal selection can generate co-evolutionary dynamics in which adaptations in one species alter the selective environment experienced by the other (Abrams 2000; Schmitz 2017). Classic examples include the escalating toxicity of rough-skinned newts and resistance of their garter snake predators, as well as the reciprocal evolution of echolocation in bats and ultrasonic hearing and evasive behaviours in their moth prey (Brodie and Brodie 1990; Brodie et al. 2005; Corcoran et al. 2009; Feldman et al. 2009; Barber et al. 2015). These predator-prey dynamics may be particularly complex during intraguild predation, in which competing species that exploit the same resources also interact as predator and prey (Polis 1981; Polis et al. 1989; Polis and Holt 1992). Intraguild predation therefore combines the selective pressures of predation and resource competition, potentially favouring improved predatory efficiency in the predator while also selecting for traits that reduce vulnerability in its prey.

Despite the importance of predator-prey interactions in shaping ecological communities and evolutionary trajectories, the mechanisms underlying reciprocal adaptation are difficult to dissect experimentally. This is particularly true for interactions between closely associated species in natural, resource-limited environments, where predation is embedded within a broader network of competition and environmental pressures. Nematodes provide an unusually tractable system for addressing these questions. Multiple nematode species commonly coexist within the same habitats, where they occupy diverse trophic niches ranging from microbial grazing to predation (Yeates et al. 1993). Yet, despite their abundance, ecological diversity, and experimental tractability, nematode predator-prey interactions remain comparatively underexplored (Ettema 1998; Bardgett and Putten 2014; Hoogen et al. 2019).

Of the nematode species capable of predatory feeding, *Pristionchus pacificus* is the most well characterised and provides an opportunity to investigate the ecological and mechanistic basis of predation (Sommer and Lightfoot 2022). It is an omnivorous species that feeds on bacteria but also acts as a predator of other nematodes (Wilecki et al. 2015). Predation is dependent on teeth-like structures that occur in two different forms within a population. The stenostomatous (St) morph has a narrow mouth with a single tooth and feeds only on bacteria, while the eurystomatous (Eu) morph has a wider mouth that contains two teeth and is capable of both bacterial feeding as well as predation on other nematodes (Ragsdale et al. 2013). In the predatory morph, contact with another nematode can trigger an attack. This action can be driven by nutritional demands but may also be an aggressive response to remove potential competitors (Quach and Chalasani 2022). The attack decision is modulated by the nematode noradrenergic system that switches the organism’s behavioural states between aggressive and docile modalities (Eren et al. 2026). Prey contact during aggressive bouts is sensed through both mechanosensory and chemosensory receptors, and results in the puncturing of the outer cuticle layer that is usually fatal for the victim (Moreno et al. 2019; Roca et al. 2026). The coordinated actions of the teeth and pharyngeal musculature are required for this activity and this is regulated through the neuromodulator serotonin (Okumura et al. 2017; Ishita et al. 2021). Moreover, predatory aggression is not indiscriminate, as *P. pacificus* has evolved a kin-recognition system that allows close relatives to be identified, and kin contact does not result in attacks (Lightfoot et al. 2019, 2021; Hiramatsu and Lightfoot 2023). Importantly, the dissection of the molecular pathways involved in predation has mostly been achieved through investigating interactions between *P. pacificus* with the distantly related model nematode species, *Caenorhabditis elegans,* although these species often do not share the same ecological niche (Félix and Duveau 2012).

While much still remains unknown regarding the ecology of *P. pacificus*, ongoing worldwide sampling efforts continue to identify new *P. pacificus* isolates as well as uncover previously unknown *Pristionchus* species (Mayer et al. 2007; Rödelsperger et al. 2014, 2018). As such, more of the complex ecology of these organisms is being revealed. *Pristionchus* species are reliably found in close association with scarab beetles with which they share a necromenic association (Herrmann et al. 2007; Kanzaki et al. 2021; Sommer 2025). This is a specific lifestyle whereby these nematodes reside on a beetle host as arrested, non-feeding, stress-resistant individuals called dauer larvae, the evolutionarily conserved dispersal stage of free-living nematodes (Sommer 2025). Many of these nematodes stay associated with their beetle host until insect death occurs via natural causes. This event results in the establishment of a stable microbiome on the decaying beetle carcass and *Pristionchus* species exploit this habitat in order to develop and propagate (Meyer et al. 2017). Within this ecosystem, they display a biphasic boom-and-bust growth and dispersal strategy that, alongside a robust ability to sense insect pheromones, likely facilitates their capacity to locate and colonise new hosts (Hong and Sommer 2006; Hong et al. 2008; Renahan et al. 2021). Moreover, beetle hosts harbour not only *Pristionchus* but also other nematode species, and additionally, other soil-dwelling nematodes are likely attracted to the microbial bloom on the decaying insect carcass (Giblin-Davis et al. 2013; Koneru et al. 2016). Therefore, the resultant microhabitat is a species-rich environment consisting of many bacterial, fungal, and nematode species interacting and competing (Meyer et al. 2017). Importantly, most wild *P. pacificus* isolates exhibit the predatory Eu mouth form (Ragsdale et al. 2013), and this morphology can be rapidly induced by specific environmental cues (Bento et al. 2010; Bose et al. 2012; Werner et al. 2017; Levis and Ragsdale 2025). These features suggest that predatory aggression likely forms a central component of their behavioural interactions on the beetle carcass, although kin-recognition mechanisms ensure that such aggression is directed toward non-relatives and other species within this ecological niche (Lightfoot et al. 2019, 2021; Hiramatsu and Lightfoot 2023).

Here, we identify a naturally co-occurring nematode species that shares the necromenic niche with *P. pacificus.* Furthermore, we establish this pairing as a system for studying unidirectional intraguild predation, in which a species directly competes with its predator for the same ephemeral resources. This may impose reciprocal selection, potentially favouring predator adaptations that increase predatory efficiency, and prey traits that either reduce susceptibility to attack or optimize resource competition efficiency. Accordingly, by using isolates collected together in the field, we ask whether *P. pacificus* preys upon this naturally co-occurring species, and if so, whether predation efficiency differs from that observed with the widely used but non-co-occurring laboratory prey species *C. elegans*. We further ask whether this natural prey displays morphological or life-history traits consistent with an evolved response to predation pressure from a natural intraguild predator. By addressing these questions, this study establishes a genetically tractable system in which the reciprocal selective pressures of predation and resource competition can be dissected using ecologically relevant, naturally co-occurring species.

## Materials and Methods

### Specimen collection

Live beetle specimens belonging to the genus *Adoretus* (Dejean, 1833) were collected by hand from Parc du Colorado, La Réunion (covering area around latitude: 20°54’43.6"S, longitude: 55°25’12.4"E) (Fig. 1A, B). They were later sacrificed on nematode growth medium (NGM) plates seeded with *Escherichia coli* OP50 at the Max Planck Institute for Biology Tübingen. A mixture of *E. coli* and native beetle carcass microbes served as a food signal for nematode dauer emergence (Kanzaki et al. 2021).

**Figure 1.**
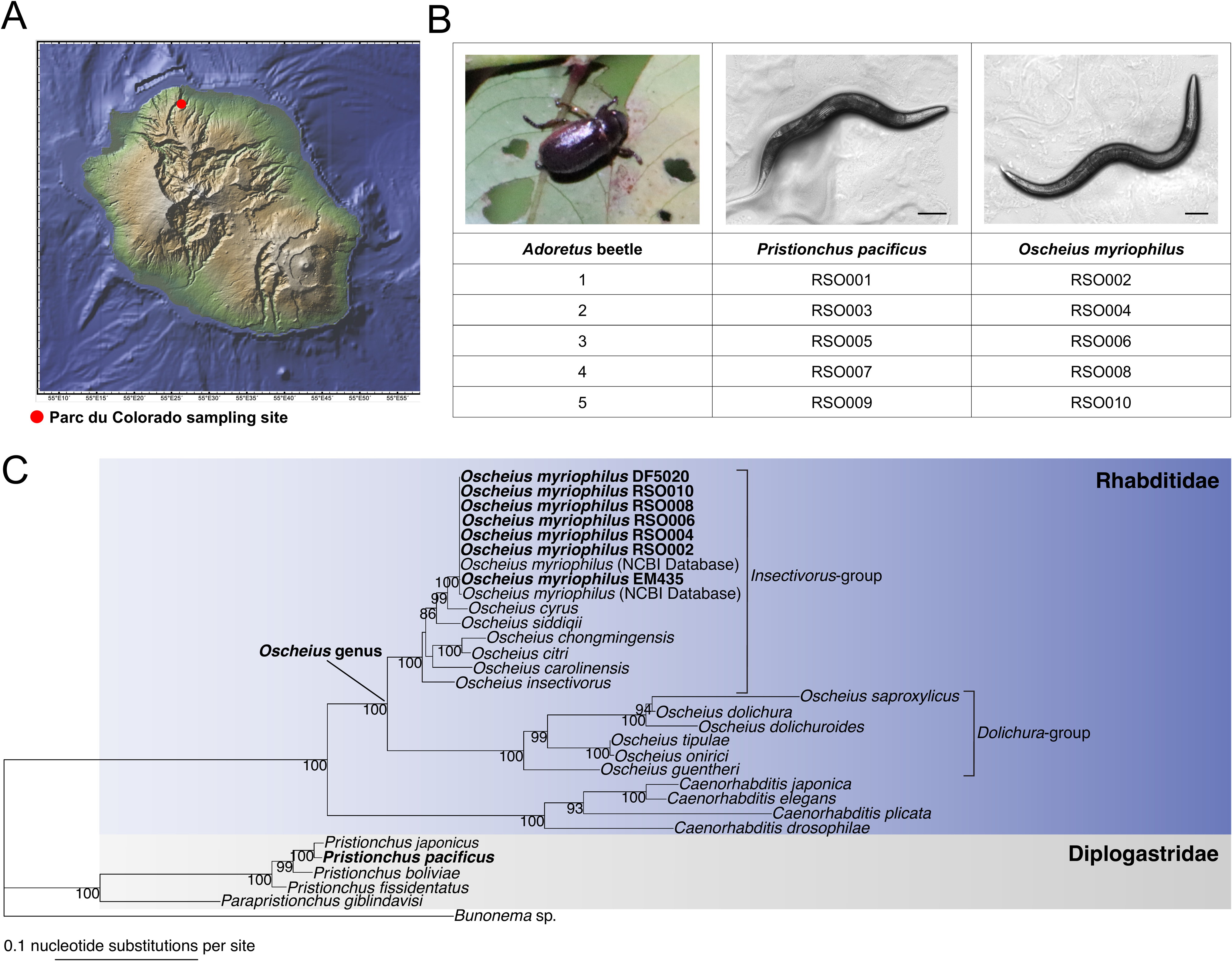
Pristionchus pacificus and Oscheius myriophilus consistently cohabitate on Adoretus beetles. (A) Map of La Réunion Island showing the approximate location of the Parc du Colorado sampling site where beetles from the genus *Adoretus* were collected. (B) Corresponding *P. pacificus* and *O. myriophilus* isolates collected from each *Adoretus* beetle, along with representative images of each species. Scale bars indicate 100 μm. (C) Maximum-likelihood tree showing relatedness among the wild *O. myriophilus* isolates collected in this study to the previous *O. myriophilus* strains EM435 and DF5020, as well as other nematodes within the family Rhabditidae, and *P. pacificus* within the family Diplogastridae. Scale bar indicates 0.1 nucleotide substitutions per site.

General body and pharyngeal morphology were used to identify the emerging *P. pacificus* and *Oscheius* nematodes (Herrmann et al. 2010; Harry et al. 2022) followed by molecular identification (see below). Each isogenic nematode strain was established from a single gravid hermaphrodite, and frozen within ten generations or less for long-term storage to avoid domestication (O’Connell 2022).

### Nematode husbandry

All nematodes were maintained on standard NGM plates on a bacterial diet of *E. coli* OP50 at 20°C (Sommer et al. 1996).

### O. myriophilus molecular identification and phylogenetic reconstruction

Two rDNA regions of the *Oscheius* isolates originating from La Réunion Island and those deposited at the *Caenorhabditis* Genetics Center (CGC) were sequenced including the near full-length small subunit (SSU) and the D2-D3 region of large subunit (LSU). The primer pairs used were SSU988F (5’-CTCAAAGATTAAGCCATGC-3’) and SSU2646R (5’-GCTACCTTGTTACGACTTTT-3’) and D2A (5’-ACAAGTACCGTGAGGGAAAGT-3’) and D3B (5’-TCGGAAGGAACCAGCTACTA-3’) for the SSU and LSU regions, respectively (Kanzaki and Giblin-Davis 2025). The sequences were used for BLASTN search in the NCBI database with *O. myriophilus* being the top hits (>99% identical).

To reconstruct the *O. myriophilus* phylogenetic relationships, the SSU and D2-D3 region of LSU sequences of *Oscheius*, *Caenorhabditis*, *Pristionchus*, *Parapristionchus* and *Bunonema* genera available on the NCBI database were concatenated and aligned using the MUSCLE program (version 3.8.31) (Table S2) (Edgar 2004). The resultant multiple sequence alignment was then used to reconstruct a maximum-likelihood tree using the phangorn package in R (version 4.3.1; model = “GTR+G(4)”, optNni = TRUE) (Schliep 2010). The best tree model was determined using the pml_bb function. Representing a known outgroup of the families Diplogastridae and Rhabditidae. *Bunonema* sp. was used for tree rooting (Qing et al. 2024; Rödelsperger et al. 2024). 100 bootstrap replicates were set for this rooted tree and nodes with the support values more than 50 were visualized on the generated tree (Figure 1C).

### P. pacificus whole genome sequencing and phylogenetic reconstruction

Genomic DNA of *P. pacificus* natural isolates co-existing with *O. myriophilus* was extracted using the Monarch^®^ Genomic DNA Purification Kit. The whole genome sequencing (WGS) was done by Novogene. The phylogenetic reconstruction was carried out following the protocol in Rödelsperger *et al*. (Rödelsperger et al. 2017). Briefly, the new WGS data were aligned against the reference *P. pacificus* PS312 genome (version El Paco 3, pristionchus.org) with the software BWA (version 0.7.17-r1188, with option mem) (Li and Durbin 2010). The resulting alignments were then genotyped by the SAMtools mpileup (version 0.1.18 r982:295) and BCFtools (version 0.1.17-dec r973:277) at a set of candidate positions that were shown to be variable across different *P. pacificus* natural isolates Rödelsperger, C. *et al*. (Rödelsperger et al. 2014, 2017; McGaughran et al. 2016; Danecek et al. 2021). Variable positions that could be genotyped in all strains with a minimum quality score of 20 were concatenated into an artificial multiple sequence alignment. This variant information was used to calculate a neighbour-joining tree based on Hamming distances between strains with the phangorn package in R (version 4.3.1) (Fig. S1) (Schliep 2010).

### La Réunion Map

The La Réunion Island map was generated using the software GeoMapApp (Ryan et al. 2009).

### Mouth-form assays

To determine whether the wild *P. pacificus* isolates collected in this study are capable of predation, individual animals were examined and classified as having either the predatory Eu or the non-predatory St morphotype (Ragsdale et al. 2013). This mouth-form analysis of *P. pacificus* isolates was performed by picking adult worms into 20 mM sodium azide to immobilize worms, mixed with M9 on a 24 x 24 mm glass slide. Given that the mouth form and teeth are fully developed and fixed in adult worms, the addition of sodium azide and resulting cessation of thrashing, aids in mouth-form identification without impacting mouth-form fate.

Following the addition of a cover glass, the mouth form of each animal was examined at 60x/oil immersion on an inverted microscope (Zeiss Axiovert 200). Animals were characterized as stenostomatous or eurystomatous based on their mouth width and the presence of either one or two teeth (Figure 2A, B). Two replicates, containing approximately 50 worms each, were performed on separate days for every isolate. Animals examined for mouth form were discarded and were not used for any additional experiments.

**Figure 2.**
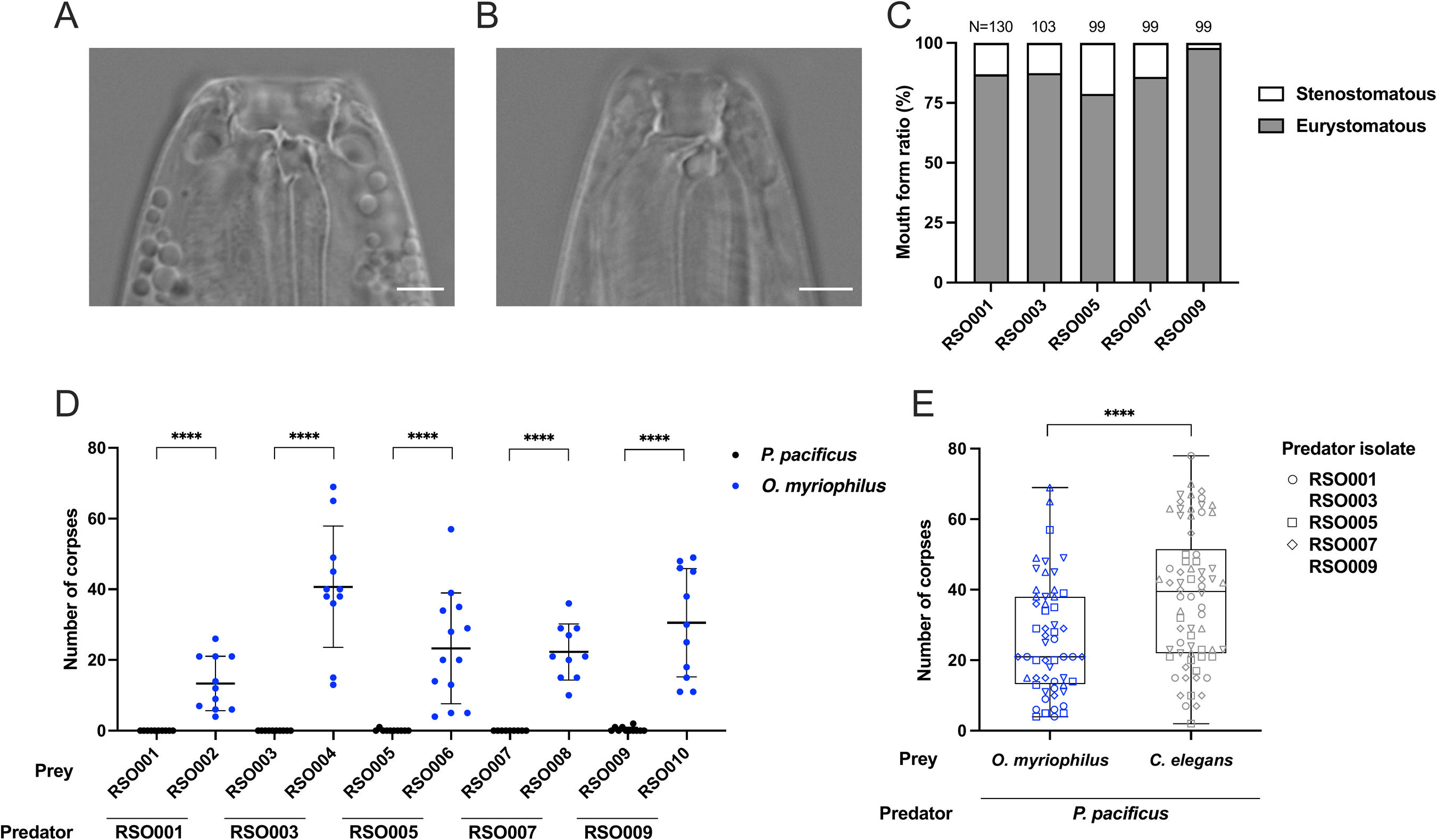
Naturally co-occurring *P. pacificus* and *O. myriophilus* demonstrate a predator-prey dynamic. (A) Representative image of an animal with the predatory eurystomatous mouth form. Scale bar indicates 5μm. (B) Representative image of the nonpredatory stenostomatous mouth form. Scale bar indicates 5μm. (C) Percent occurrence of the eurystomatous and stenostomatous mouth forms in the wild *P. pacificus* isolates collected in this study. N ≥ 99 animals per isolate. (D) Number of corpses present after five *P. pacificus* adult predators were exposed to larval prey that is either self-progeny or their naturally co-occurring *O. myriophilus* isolate for two hours. Data points represent individual trials (N ≥ 10) with mean ± SD shown. **** *p* < 0.0001, Mann-Whitney *U* test. (E) Combined *O. myriophilus* corpse counts from panel D compared to predation levels on *C. elegans* reveals that *P. pacificus* demonstrates higher levels of predation on *C. elegans* than *O. myriophilus*. Data points represent individual trials. Box plots shown with the first to third quartiles indicated by the box, the median by the center line, and the minimum and maximum values by the whiskers. **** *p* < 0.0001, Wald *z*-test.

### Lifespan and fertility assays

To investigate the life history traits of the co-occurring isolate pairs, lifespan and fertility were assessed in tandem. On day 0, early J4 hermaphrodites were selected based on their vulval morphology and picked onto individual NGM 6 cm plates seeded with *E. coli* OP50. The following day was counted as day one of adulthood. Hermaphrodites were transferred to fresh plates as necessary (every day during early fertility or every other day as fertility declined with age) while still fertile to avoid confounding parent and progeny. Eggs on progeny plates were allowed to hatch and then the number of live offspring were quantified and recorded as the total number of lifetime progeny per individual worm. While fertile, hermaphrodites were examined daily for internal hatching of progeny and, if present, the initial date of occurrence was recorded. Each parent was then followed until the date of their death was determined based on a complete lack of response to a harsh touch. The number of days of adulthood was recorded and dead animals were maintained as long as necessary for all live progeny to escape.

### Predatory corpse assay

To determine whether *P. pacificus* predators prey on their co-occurring *O. myriophilus* isolate, corpse assays were performed as in (Wilecki et al. 2015). For this assay, large quantities of each isolate to be utilized as prey were cultivated on standard NGM plates seeded with *E. coli* OP50. The plates were allowed to grow until the population had just depleted the bacterial food source and then the freshly starved plates were gently rinsed with sterile M9. The M9/worm solution was then poured through two 20 μm filters stacked on top of each other to separate adults from the J1 and J2 larvae that are small enough to flow through the filters. The adults and larger worms captured on top of the filters were discarded, and the solution containing the larvae was spun at 2000 rpm for one minute to collect the larvae at the bottom of the tube. The supernatant was discarded and two more wash and spin cycles were performed with M9 to aid in removal of any remaining bacteria. Subsequently, 2 μl of the larval prey pellet was then pipetted to an unseeded 6 cm NGM plate and the larval prey were allowed to disperse for approximately two hours. This results in an NGM assay plate that is saturated with larval prey and devoid of any bacterial food source. For *P. pacificus* or *C. elegans* prey, pipetting a single volume of prey to the centre of the plate is sufficient to allow for adequate dispersal. However, we noted that *O. myriophilus* prey does not disperse as readily. Thus, *O. myriophilus* prey were pipetted to two offset regions of the plate to aid in dispersal (1 μl in each region). Separately, well-fed, young adult predators were picked to unseeded plates and left off food for one hour prior to the start of the assay to remove residual bacteria adhering to their body and induce mild starvation. Finally, five adult predators were placed on each prey-saturated assay plate for a period of two hours.

During the two-hour period, predators are able to prey on the larval prey, and successful predatory events leave behind a visible corpse. At the end of the two-hour period, the predators were removed and the number of corpses were counted manually using a stereo microscope (Zeiss Stemi 508). The total number of corpses generated by the five predators on each assay plate was recorded as a measure of the level of predation that occurred.

### Killing efficiency assay

To assess the effectiveness of *P. pacificus* predatory bites on *O. myriophilus* prey of different developmental stages, we isolated two different developmental stages of RSO006 to be used as prey, namely J1/J2 larvae and J4/young adult animals. The *O. myriophilus* J1/J2 early-stage larvae were collected as described above for corpse assays and 1 μl of larvae was pipetted onto a 6 cm unseeded plate. A lower prey density than for a corpse assay was utilized to aid in visual examination of individual interactions between the predator and prey. The larvae were left to disperse for two hours and then >10 young adult *P. pacificus* RSO005 predators were placed onto the *O. myriophilus* larval plate. Individual predators were observed interacting with *O. myriophilus* larval prey using a Zeiss Stemi 508 until they had attempted a total of ten bites.

Subsequently, the number of successful ‘kills’ out of the ten total ‘bites’ was recorded. Interactions were scored as ‘bites’ if the predator successfully grasped the larval prey and prevented their locomotion. ‘Kills’ were scored if a bite resulted in the penetration of the cuticle and subsequent death of the larvae. To assess *P. pacificus* predator killing efficiency against *O. myriophilus* J4/young adults, we picked ∼300 RSO006 J4/young adults to a 6 cm unseeded plate. The J4/young adult stage was determined by the presence of the classic vulva morphology during this developmental period. Greater than 10 young adult *P. pacificus* RSO005 predators were subsequently placed onto the *O. myriophilus* J4/young adult plate, and the number of ‘bites’ and ‘kills’ were assessed as described above. A total of 100 predatory interactions were observed across 10 independent predators for each prey type.

### O. myriophilus matricide on natural beetle associated bacteria

The influence of bacterial diet on the reproductive mode of *O. myriophilus* was assessed by growing *O. myriophilus* RSO006 on four different bacterial species that have previously been found naturally co-occurring on beetle carcasses found on La Réunion Island (Akduman et al. 2018). These were *Pseudomonas fluorescens* LRB26, *Kurthia gibsonii* LRB56, *Comamonas thiooxydan* LRB28, and *Providencia rettgeri* LRB44 (Athanasouli et al. 2025). A starter culture of each of these bacteria was grown overnight at 25 °C and 200 μl was then seeded onto standard NGM plates. These were grown for 24 h at room temperature before *O. myriophilus* RSO006 eggs were added to each plate. *O. myriophilus* were then allowed to hatch and grow for 4-5 days on the bacterial plates before being examined for matricidal hatching, as evidenced by the presence of internally hatched progeny and parental death (Zeiss Axio Zoom V16).

### Statistics

Unless otherwise stated, statistical analyses were performed using GraphPad Prism (version 11.0.2). As many individual isolate datasets did not satisfy the assumptions of parametric tests, as indicated by the Shapiro-Wilk test for normality, non-parametric statistical tests were utilized for pairwise comparisons of corpse counts, lifespan, and fertility between each of the co-occurring predator-prey isolate pairs. These pairwise comparisons were performed using two-sided Mann-Whitney *U* tests. Categorical data (bagged vs. oviposition only) were analysed with two-sided Fisher’s exact tests. Exact sample sizes for each experiment are reported in the respective figure legends and statistical significance was defined as *p* < 0.05. The experimental unit was defined according to the specific design of each assay. For corpse assays, each assay plate containing five predators represented one independent biological replicate. For mouth-form assessment, as well as lifespan and fertility experiments, each individual animal represented one biological replicate. For the killing efficiency assay, a single predator represented one biological replicate. No single animal was ever used for multiple assays.

In Figure 2, corpse counts for *C. elegans* and *O. myriophilus* were analysed using a negative binomial generalized linear mixed-effects model to account for overdispersion in the count data and variation among predator isolates. This analysis was performed in R (version 4.4.2) using the glmmTMB package (version 1.1.15.2), with prey species included as a fixed effect and predator isolate included as a random intercept. A model including a random slope for prey species by predator isolate was compared with the random-intercept model using a likelihood ratio test but this did not significantly improve the model fit (χ² = 2.95, df = 2, *p* = 0.2284), and thus the simpler random-intercept model was retained. Estimated marginal means and pairwise comparisons between prey species were calculated using the emmeans package (version 2.0.2), with significance assessed using a Wald *z*-test. The contrast was back-transformed to the response scale and reported as a ratio of expected corpse counts.

To determine whether the onset of ovoviviparity was associated with subsequent mortality in *O. myriophilus* in Figure 4, we fit a Cox proportional hazards model that included ovoviviparity as a time-dependent covariate and isolate as a fixed effect. Individual worms contributed observation time to the non-ovoviviparous state until internal hatching of progeny was first observed and to the ovoviviparous state thereafter. The proportional hazards assumption was evaluated using Schoenfeld residuals. As a modest departure from this assumption was detected for ovoviviparity, a second Cox model incorporating a time-varying coefficient for ovoviviparity was fit as a sensitivity analysis, yielding the same qualitative conclusion and indicating that the primary finding is robust to the modest violation of this assumption. Statistical analyses were performed in R (version 4.4.2) using the survival package (version 3.7-0). Significance of individual model coefficients was assessed using Wald *z*-tests and the 95% confidence interval was calculated from the estimated regression coefficients.

Killing efficiency data in Figure 4 were analysed in R (version 4.4.2) using a binomial generalized linear model with prey developmental stage as a fixed effect. This analysis was utilized rather than a Fisher’s exact test because each predator contributed 10 observations to the data. Each predator was treated as one biological replicate, and a binomial error distribution was used because the response variable was the number of successes (kills) out of a fixed number of encounters with (ten bites) individual prey. Model dispersion was assessed using the Pearson dispersion statistic (0.80), indicating no evidence of overdispersion, and significance was assessed with a likelihood ratio test.

## Results

### Identification and isolation of nematode predator-prey systems

To better understand the behavioural interactions between *P. pacificus* and its natural prey species, we set out to identify and isolate nematodes that naturally co-occur with these facultative predators. Given that *P. pacificus* is reliably, although not exclusively, found on beetle hosts (Félix et al. 2018), we conducted field experiments on La Réunion Island to capture beetles of the family Scarabaeidae that may harbour *P. pacificus* nematodes and their prey (**Fig. 1A**). On La Réunion Island, beetles have been previously demonstrated to have a *P. pacificus* infestation rate of over 90% (Herrmann et al. 2010). At the Parc du Colorado sampling site on La Réunion Island (covering area around latitude: 20°54’43.6"S, longitude: 55°25’12.4"E), we captured numerous beetles belonging to the genus *Adoretus* that were infested with several nematode species (**Fig. 1A, B; Supp Table 1**). Initial morphological analysis indicated these nematode species primarily belonged to the family Rhabditidae due to the presence of a pharyngeal grinder, and the family Diplogastridae, identified by their characteristic pharyngeal shape and structure (Harry et al. 2022). We focused on five *Adoretus* isolates that each harboured both a single diplogastrid and a single rhabditid individual. As the isolated specimens were hermaphroditic species, we used the individual nematodes to initiate an isogenic master culture of each that was then maintained as an independent line and frozen stock (**Fig. 1B**). To precisely determine the species composition harboured on these five beetles, we performed molecular genotyping by whole genome sequencing the diplogastrids, and sequencing the rhabditids using universal nematode molecular markers followed by phylogenetic analysis. All diplogastrid isolates were *P. pacificus* and belong to clade C1 with other *P. pacificus* specimens isolated from La Réunion Island (Rödelsperger et al. 2017) (**Supp Fig. 1**). Alongside these, the rhabditids were all identified as specimens of *Oscheius myriophilus* based on the near full-length small subunit

(SSU) and the D2-D3 region of large subunit (LSU) sequences (Poinar 1986) (**Fig. 1C**). The monophyletic genus *Oscheius* comprises two groups (clades), *Insectivorus*- and *Dolichura*-groups (Sudhaus and Hooper 1994), and consists of many free-living members including the well-established satellite model species *Oscheius tipulae* (Ahn and Winter 2006; Felix 2006; Rosa et al. 2020; Dockendorff et al. 2022). Members of this genus, including *O. myriophilus*, are frequently associated with insects, and this species has previously been recovered from a broad range of insect hosts. (Poinar 1986; Demir et al. 2017; Ghavamabad et al. 2021). Thus, across five *Adoretus* beetle hosts, we recovered distinct *P. pacificus* strains, each co-occurring with an *O. myriophilus* isolate. These repeated associations identify an ecologically relevant species pair and provide a natural context in which to investigate their ecological interactions.

### P. pacificus are predators of O. myriophilus

Having established the ecological association between these nematodes, we next examined whether these pairs constitute bona fide predator–prey relationships. We first assessed the mouth morphology ratios in these newly acquired wild *P. pacificus* strains to determine their predatory capability. In *P. pacificus*, only the Eu morph exhibits aggressive predatory behaviour and this behaviour is entirely absent in the St morph (**Fig. 2A, B**) (Wilecki et al. 2015). As most wild isolates characterised to date show a strong bias toward the Eu fate (Ragsdale et al. 2013), we assessed all five of our beetle-associated *P. pacificus* strains for the presence of the Eu mouth type. In all strains we identified a similar Eu bias confirming the ability of these isolates to develop the predatory-capable morphotype (**Fig. 2C**). Although mouth-form ratio is influenced by environmental factors (Werner et al. 2017, 2018; Lenuzzi et al. 2023; Piskobulu et al. 2025), and thus could be influenced by laboratory conditions, the observed bias toward the predatory Eu morph is consistent with previous findings in other wild *P. pacificus* lineages (Ragsdale et al. 2013).

With the predominance of the predatory morph confirmed under laboratory conditions, we next investigated predatory interactions between *P. pacificus* and their co-occurring *O. myriophilus* isolate. To this end, we conducted predatory “corpse assays” whereby we placed several *P. pacificus* predators on an assay plate with an abundance of *O. myriophilus* larval prey. After two hours, the number of corpses generated by predatory events were counted indicating the amount of predatory activity (Wilecki et al. 2015; Lightfoot et al. 2016). We found that all of the wild *P. pacificus* isolates do indeed prey upon their naturally co-occurring *O. myriophilus* isolate, confirming the occurrence of a predator-prey dynamic between the species (**Fig. 2D**). In comparison, corpse assays utilizing self-progeny as larval prey revealed that all *P. pacificus* isolates robustly avoid killing their own kin (**Fig. 2D**), consistent with the existence of a kin-recognition system in this species (Lightfoot et al. 2019, 2021). If cohabitation imposes selective pressures on predator and prey, then we might expect predation efficiency on *O. myriophilus* to differ from predation efficiency on other non-cohabitating prey species. To investigate this, we performed corpse assays using *C. elegans* prey, which has commonly been utilized as prey in previous publications but cohabitates infrequently with *P. pacificus* (Félix and Duveau 2012), and compared these data to the pooled *O. myriophilus* corpse counts. A negative binomial generalized linear mixed model with predator isolate as a random intercept revealed that *P. pacificus* predators produced significantly more corpses when exposed to *C. elegans* prey, and that corpse counts on *C. elegans* were 1.50-fold higher than on *O. myriophilus* (95% CI 1.23– 1.83; Wald *z* = 3.97, *p* < 0.0001) (**Fig. 2E)**. Together, these findings establish intraguild predation between *P. pacificus* and *O. myriophilus,* and reveal that predation efficiency varies when predators are exposed to intraguild vs. extraguild prey.

### Distinct life history traits are evident in co-occurring predator-prey species

After verifying the ecological and behavioural association between these nematodes, we next examined their life-history traits for evidence of any potential adaptations linked to their respective trophic strategies and reciprocal interactions. These traits included their development, fertility, and longevity.

To assess the development and longevity of our co-occurring predator-prey species, we followed individual animals from young adults (one day post J4) until their death. The adult lifespan of *P. pacificus* varied between isolates with the shortest lifespan of 12.6 ± 2.9 days observed in RSO009 and the longest lifespan of 33.3 ± 9.0 days observed in RSO005 (mean ± SD) (**Fig. 3A**). We noted that there was less variability in longevity among the *O. myriophilus* isolates, which had adult lifespans ranging from only 8.0 ± 4.6 days in RSO002 to 10.1 ± 3.3 days in RSO004 **(Fig. 3B).** The mean adult lifespan of all the wild *P. pacificus* isolates combined was 23.0 ± 10.7 days compared to only 9.0 ± 4.3 days for the *O. myriophilus* isolates (**Fig. 3C)**. Additionally, we noted that the *O. myriophilus* isolates have significantly shorter adult lifespans than their co-occurring *P. pacificus* isolates, but that the magnitude of this effect varied among isolate pairs (**Fig. 3C**). While the majority of *P. pacificus* individuals produced offspring through oviposition, in all the *O. myriophilus* specimens we instead observed a mixed reproductive mode. This began with the oviposition of a few eggs followed by a consistent bagging phenotype, where eggs are retained and hatch internally (**Fig. 3D-F, Supp video 1)**. Death of the hermaphrodite often occurred within a few days after progeny began hatching internally. To determine whether the onset of bagging was associated with subsequent mortality, we modelled internal hatching of progeny as a time-dependent covariate in a Cox proportional hazards model while controlling for isolate. This model revealed that internal hatching of progeny was associated with a significantly increased hazard of death (HR = 10.39, 95% CI = 3.31–32.59, *p* < 0.0001) (**Fig. 3B**). This suggests that this evolved retention of embryos is correlated with a shortened lifespan, and is likely a form of matricide resulting from trauma incurred by retained progeny that are growing rapidly, thrashing, and consuming internal tissues. Furthermore, embryo retention and matricide were also observed when *O. myriophilus* isolates were grown on native beetle-associated bacteria (Akduman et al. 2018, 2020; Athanasouli et al. 2025), demonstrating that this phenotype is not specific to standard laboratory nematode culture conditions. (**Supp Fig. 2**). To investigate the origins of the mixed reproductive mode further, we assessed two additional strains of *O. myriophilus*, EM435 and DF5020, which were obtained from the *Caenorhabditis* Genetics Center (CGC). *O. myriophilus* EM435 was isolated from a compost pile in Brooklyn, New York and *O. myriophilus* DF5020 was isolated from the millipede *Oxidus gracilis* in Azusa, California (Poinar 1986). In both strains, we again observed matricidal hatching, although this phenotype was not fully penetrant in strain EM435 (**Supp Fig. 3**). Notably, many EM435 individuals were sterile and alongside this these animals exhibited extended lifespans, further supporting a link between internal hatching and matricide in *O. myriophilus*. Together, these findings suggest that natural variation of this reproductive mechanism exists within the species. Importantly, this reproductive strategy resulted in many *O. myriophilus* larvae breaking free of their mother’s carcass at a much later developmental stage in all strains examined, sometimes even as fertile adults (**Fig. 3F, Supp Video 2**), indicating that the internal contents of the mother’s body must serve as a nutrient source for the developing larvae.

**Figure 3.**
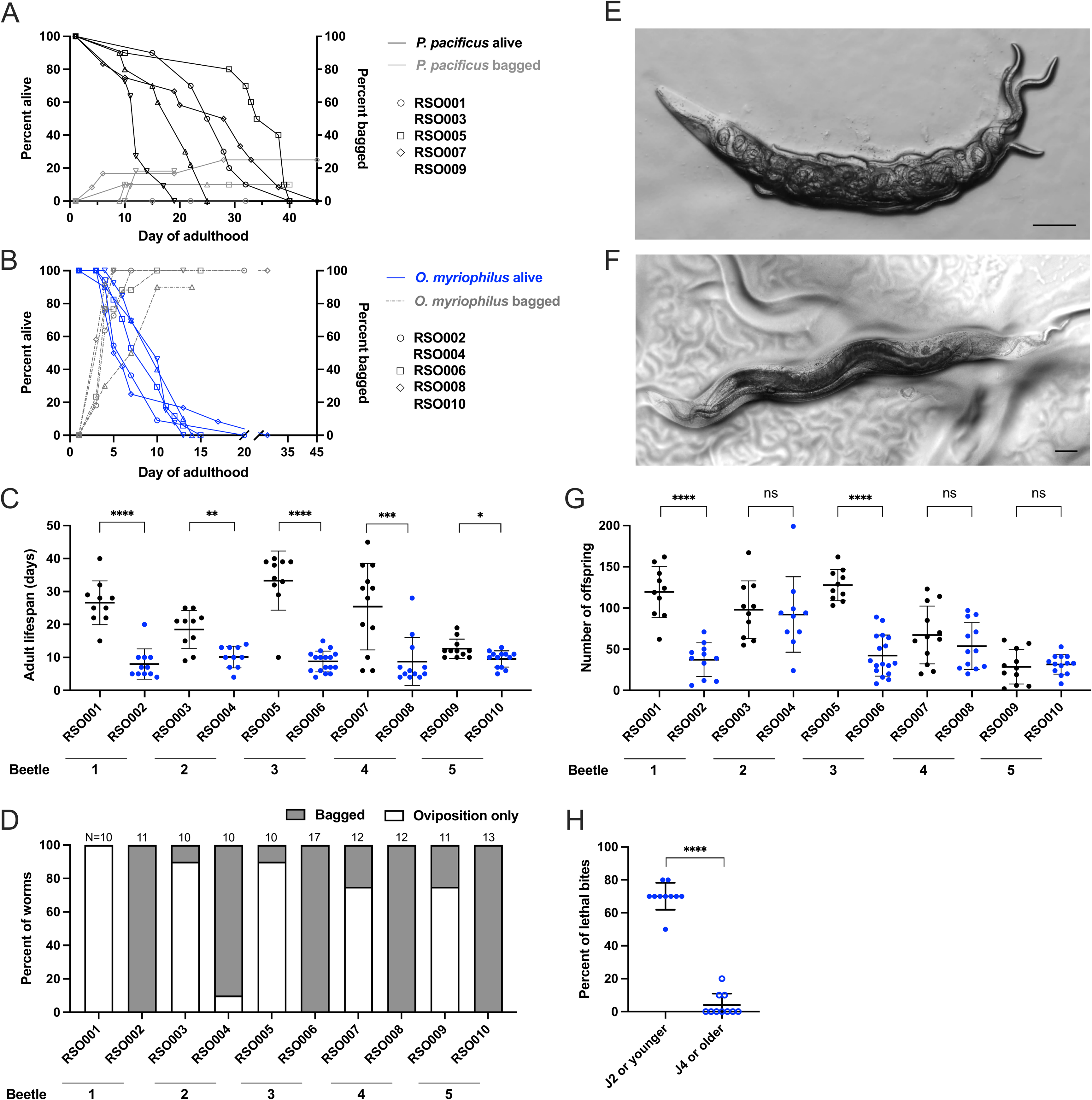
Matricidal hatching is a fixed trait in *O. myriophilus* isolates. (A-B) Adult lifespan plotted against the onset of hatching of internal progeny for *P. pacificus* isolates (A), and *O. myriophilus* isolates (B). The onset of internal egg hatching and bagging is associated with a significantly increased hazard of death in *O. myriophilus*. *p* < 0.001, Wald *z*-test. Day 0 is when animals are picked as early J4 larvae. N ≥ 10 animals per isolate. (C) *P. pacificus* isolates have a longer adult lifespan than their co-occurring *O. myriophilus* isolates. Data points indicate the adult lifespan of one individual animal with mean ± SD shown. (D) Nearly 100% of *O. myriophilus* individuals from each isolate show matricidal hatching, in contrast to *P. pacificus* isolates, where internal hatching of progeny occurs significantly less frequently. *p* < 0.0001 for all pairwise comparisons between co-occurring isolates, Fisher’s exact test. (E-F) Representative images of internally hatched progeny at various developmental stages in *O. myriophilus,* spanning from early-stage larvae (E) to gravid adults (F). Scale bars indicate 100 μm. (G) Two *P. pacificus* isolates have higher levels of fertility than their cohabitating *O. myriophilus* isolate. Data points indicate the number of live offspring from one individual with mean ± SD shown. (H) Percent of *O. myriophilus* RSO006 individuals, either early-stage larvae (J1 or J2) or animals J4 or older, that suffered a lethal bite from an adult *P. pacificus* RSO005 predator. Older animals are significantly more likely to survive a predatory attack. **** *p* < 0.0001, likelihood ratio test. Individual data points represent the percent of prey killed out of ten total prey that were bitten by a single predator. Mean ± SD of N = 10 predators per prey type shown. (C, G) ns = not significant, * *p* < 0.05, ** *p* < 0.01, *** *p* < 0.001, **** *p* < 0.0001, Mann–Whitney *U* test.

Given the consistent mixed reproductive mode and matricide observed in *O. myriophilus*, we next assessed the fertility of both *P. pacificus* and their co-occurring *O. myriophilus* by quantifying the number of live offspring produced. We found a large variation of within species fertility with *P. pacificus* RSO005 averaging the most offspring (127.7 ± 18.9 larvae) compared to RSO009 with the least (28.6 ± 20.8 larvae) (mean ± SD) (**Fig. 3G**). A similar phenomenon was observed within *O. myriophilus* with the greatest difference observed between RSO004 and RSO010 strains (92.1 ± 45.8 and 31.46 ± 11.8 larvae, respectively). Interestingly, we noted that two of the *P. pacificus* isolates had significantly more offspring than their co-occurring *O. myriophilus* isolates, while the other three isolate pairs had similar numbers of offspring (**Fig. 3G**). This suggests that the reproductive mode observed in the *O. myriophilus* isolates likely has only a negligible effect on the total number of live offspring produced. Taken together, these findings raise the possibility that while the lengthened lifespan and higher numbers of progeny observed in certain *P. pacificus* isolates may contribute a fitness advantage, the fixation of matricidal hatching among the *O. myriophilus* isolates may represent an alternative adaptive strategy within the shared beetle carcass environment.

Given that the observed reproductive strategy in *O. myriophilus* resulted in some progeny reaching later developmental stages before breaking free of their mother’s cuticle, we hypothesized that this delayed exposure to the predatory environment may provide a survival advantage. To investigate this, we initially aimed to quantify the specific ages of progeny when they escaped from the mother’s carcass to determine what percent escaped at later developmental stages. Unfortunately, these efforts were confounded by the fact that larvae often remained within the dead carcass for multiple days, during which, bacterial contamination from the carcass accumulated on the agar plate. As a result, escaped progeny continue to grow and develop in the presence of this bacterial food source. Thus, determination of the precise developmental stage of the progeny at the time of escape was not be feasible without continuous high-resolution monitoring over multiple days. However, we regularly observed both early and later stage larvae emerging from the carcasses of all of the *O. myriophilus* isolates, but noted that there was variability between individual carcasses (**Fig 3E-F).** Finally, as we observed some animals almost completing their development within their mother’s carcass, we assessed the impact of developmental stage on avoiding fatal predatory interactions to determine if late-stage larvae might have a survival advantage. We directly observed the effectiveness of *P. pacificus* predatory bites on distinct *O. myriophilus* developmental stages by comparing bite effectiveness on early-stage larvae (J1 or J2) to bite effectiveness on the last larval stage (J4) or young adults. A binomial generalized linear model revealed that kill frequency differed significantly between young and old prey (likelihood ratio test: χ² = 107.82, *p* < 0.0001), with bites on J1 or J2 larvae resulting in a kill 70% of the time, whereas only 4% of bites on J4 or adults were fatal (**Fig. 3H**). Thus, *P. pacificus* bites were much less effective against later developmental stages of *O. myriophilus.* We speculate that this reduced lethality may be related to an increase in cuticle thickness across development, consistent with similar observations in other nematode species (Wilecki et al. 2015; Ichiishi et al. 2021). Therefore, while direct measurements of cuticle thickness across *O. myriophilus* developmental stages are needed to confirm this hypothesis, emerging from the mother’s carcass at a more advanced developmental stage may provide protection against predation.

## Discussion

*P. pacificus* represents the best-characterised predatory nematode species, yet interactions with its natural prey have remained largely unexplored. Such antagonistic interactions exert strong reciprocal selection, potentially influencing both behavioural and developmental traits. Here, we identify *O. myriophilus* as a natural prey for *P. pacificus* and establish this species pairing as a tractable system for investigating intraguild predation. As both species coexist on the same transient necromenic resource and compete for this while one also preys upon the other, this system integrates predator–prey interactions with resource competition, providing a powerful framework for studying the ecological and evolutionary consequences of both processes.

Using our well-established corpse assays to explore predation behaviour between co-occurring *P. pacificus* and *O. myriophilus,* we observed evidence of abundant predator-prey interactions. Unexpectedly, we found *P. pacificus* preys less upon its intraguild prey species, *O. myriophilus,* than *C. elegans*, which it likely encounters far less often. It has previously been shown that different nematode species acquire distinct surface properties that likely reflect species-specific ecological pressures and environmental adaptations (Chauhan et al. 2017; Kotowska et al. 2025; Hiramatsu et al. 2026). Therefore, the surface of *O. myriophilus* may have acquired specific adaptations that help to mask it from the predatory *P. pacificus*. These surface modifications may represent an evolutionary countermeasure to the aggression of *P. pacificus* and would likely require continual refinement to maintain their effectiveness. Furthermore, whether *O. myriophilus* has also evolved any specific predator-avoidance behaviours that are absent or lost as a result of domestication in *C. elegans* remains an open question. Thus, establishing the avoidance response in both species as prey would further test the coevolutionary hypothesis proposed here.

In addition to reduced predation, all *O. myriophilus* isolates exhibited a specific reproductive strategy characterized by consistent internal hatching and matricide. In *C. elegans,* reproduction occurs via egg laying, and bagging, is a rare event observed mostly in response to adverse environmental conditions including starvation (Chen and Caswell-Chen 2004), aging (Pickett and Kornfeld 2013) or exposure to pathogenic bacteria (Mosser et al. 2011). These conditions are thought to cause alterations or defects in the mechanisms that modulate the egg-laying machinery itself (Pickett and Kornfeld 2013). Similarly, in *O. tipulae*, the model species of the genus *Oscheius*, matricidal hatching is a rare event (Felix 2006). However, in many nematode species this is a well-documented reproductive strategy. This includes entomopathogenic species such as those belonging to the families Steinernematidae and Heterorhabditidae (Baliadv et al. 2011). In these species, matricide is thought to facilitate transmission of their insect-pathogenic bacterial symbionts and is essential for their specific life cycle (Ciche et al. 2008; Baliadv et al. 2011). In *O. myriophilus*, internally hatched progeny are sheltered from environmental hazards and potential predation during their early development. This corresponds to when the offspring cuticle is relatively thin and has not yet developed the thicker, more protective properties characteristic of later stages (Ichiishi et al. 2021). In addition to providing a measure of protection, the dead mother’s carcass also acts as a food source for the internally developing larvae, and thus eliminates resource competition between the sheltered progeny and the intraguild predator. Furthermore, by effectively eradicating adult hermaphrodites from the population shortly after they have reproduced, matricidal hatching also reduces intraspecific competition between parents and their own offspring. Importantly, given the penetrance and ubiquity of this phenotype in the *O. myriophilus* isolates described in this study and also evident in the specimens from other locations, we hypothesize that matricidal hatching is likely the primary mechanism of reproduction and may represent an evolved strategy to increase resource competition efficiency and maximize offspring survival in the face of predation.

Intriguingly, an early period of oviposition at the onset of fertility is retained by *O. myriophilus*. As a result, early progeny hatch directly into the environment while later progeny hatch and develop within the mother. We posit that this reproductive strategy, and its inherent trade-off with the parent’s longevity, may represent a bet-hedging strategy resulting in dual offspring phenotypes that maximize fitness across unpredictable environments (Philippi and Seger 1989; Crean and Marshall 2009). Here, externally hatched larvae, though vulnerable to predation, could enter dauer diapause or disperse to locate new hosts and environments (Hong and Sommer 2006). Conversely, internally hatched progeny would be sheltered from environmental hazards and predation. By partitioning offspring between dispersal and protection strategies, *O. myriophilus* hedges against the inherent temporal variations caused by the boom- and-bust resource availability observed on its beetle carcass, as well as habitat spatial heterogeneity linked to predator presence (Renahan et al. 2021). Thus, this reproductive bet-hedging strategy maximises the possibility that at least some offspring survive regardless of external conditions.

The potential reciprocal selection we have uncovered between *P. pacificus* and *O. myriophilus* reflects a phenomenon observed across diverse predator-prey systems. Within the confined microhabitat of an insect cadaver, *P. pacificus* and *O. myriophilus* compete directly for limited resources, creating strong selective pressure for improved predatory efficiency in *P. pacificus* and the evolution of defensive counterstrategies in *O. myriophilus*. Crucially, this nematode species pair offers exceptional genetic and experimental tractability, enabling direct dissection of the molecular and developmental mechanisms that drive predator-prey intraguild conflict and co-evolutionary adaptations in the future.

## Figure Legends

**Supplemental Figure 1. Newly isolated *P. pacificus* strains belong to clade C1**

Neighbour-joining tree generated by whole genome sequencing showing the phylogenetic relationships between the *P. pacificus* isolates collected in this study (indicated by red diamonds) and other *P. pacificus* isolates collected worldwide. Letters and numbers around the edge of the phylogeny indicate *P. pacificus* clades following Rödelsperger et al (Rödelsperger et al. 2017).

Scale bar indicates 0.01 nucleotide substitutions per site.

**Supplemental Figure 2. Matricidal hatching in *O. myriophilus* occurs when grown on diverse, beetle-associated bacteria**

Representative images of dead *O. myriophilus* RSO006 adults with internally hatched progeny (white arrows) when grown on a selection of beetle-associated bacterial species, including (A) *Pseudomonas fluorescens* LRB26, (B) *Kurthia gibsonii* LRB56, (C) *Comamonas thiooxydan* LRB28, and (D) *Providencia rettgeri* LRB44. Scale bars indicate 200 μm.

**Supplemental Figure 3. Reference *O. myriophilus* strains also show matricidal hatching.**

(A-B) Adult lifespan plotted against the onset of hatching of internal progeny for *O. myriophilus* reference strains, EM435 (A), and DF5020 (B). Day 0 is when animals were picked as early J4 larvae. N ≥ 11 animals per isolate (C) Adult lifespan of *O. myriophilus* reference strains. Data points indicate the lifespan of one individual animal with mean ± SD shown. (D) Ovoviviparity occurs at varying frequencies in the two *O. myriophilus* reference strains. (E) Fertility of EM435 and DF5020. Data points indicate the number of live offspring from a single animal with mean ± SD shown.

**Movie S1. Matricidal hatching is the primary mechanism of reproduction of in *O. myriophilus*.**

**Movie S2. *O. myriophilus* can complete their development within the corpse of the mother.**

## Acknowledgements

We thank Dr Cameron Weadick (University of Exeter) and Dr Pascal Malkemper (MPI for Neurobiology of Behavior, Bonn) for their critical reading and constructive comments on the manuscript. We thank Nurit Zorn (MPI for Neurobiology of Behavior, Bonn) for strain maintenance, Dr Christian Rödelsperger (MPI for Biology, Tübingen) for bioinformatic discussions, Heike Haussmann (MPI for Biology, Tübingen) for freezing and thawing of natural nematode isolates and relevant La Réunion Island authorities for specimen collection permits. *Oscheius* strains EM435 and DF5020 were provided by the CGC, which is funded by NIH Office of Research Infrastructure Programs (P40 OD010440).

## Conflict of Interest statement

We have no conflict of interest to declare.

## Funding

This project was funded by the Deutsche Forschungsgemeinschaft (DFG, German Research Foundation) as part of the SPP 2205 – 533395631 and the Max Planck Society.

## Data Availability

All data are available in the main and supplementary files. Raw data is available upon request and WGS data have all been deposited at the European Nucleotide Archive under the project accession PRJEB108083.

## Notes

### Competing Interest Statement

The authors have declared no competing interest.

### Summary of Updates

Title has been updated Text and figures have been improved throughout

## References

Abrams PA (2000) The Evolution of Predator-Prey Interactions: Theory and Evidence. Annu Rev Ecol Syst 31:79–105. 10.1146/annurev.ecolsys.31.1.79

Ahn I-Y, Winter CE (2006) The genome of Oscheius tipulae : determination of size, complexity, and structure by DNA reassociation using fluorescent dye. Genome 49:1007–1015. 10.1139/g06-045

Akduman N, Lightfoot JW, Röseler W, et al (2020) Bacterial vitamin B12 production enhances nematode predatory behavior. Isme J 14:1494–1507. 10.1038/s41396-020-0626-2

Akduman N, Rödelsperger C, Sommer RJ (2018) Culture-based analysis of Pristionchus-associated microbiota from beetles and figs for studying nematode-bacterial interactions. Plos One 13:e0198018. 10.1371/journal.pone.0198018

Athanasouli M, Loschko T, Rödelsperger C (2025) Interspecies systems biology links bacterial metabolic pathways to nematode gene expression, chemotaxis behavior, and survival. Genome Res 35:2363–2374. 10.1101/gr.280848.125

Baliadv Y, Yoshiga T, Kondo E (2011) Development of Endotokia Matricida and Emergence of Originating Infective Juveniles of Steinernematid and Heterorhabditid Nematodes. Nematol Res (Jpn J Nematol) 31:26. 10.3725/jjn1993.31.1-2_26

Barber JR, Leavell BC, Keener AL, et al (2015) Moth tails divert bat attack: Evolution of acoustic deflection. Proc Natl Acad Sci 112:2812–2816. 10.1073/pnas.1421926112

Bardgett RD, Putten WH van der (2014) Belowground biodiversity and ecosystem functioning. Nature 515:505–511. 10.1038/nature13855

Bento G, Ogawa A, Sommer RJ (2010) Co-option of the hormone-signalling module dafachronic acid–DAF-12 in nematode evolution. Nature 466:494–497. 10.1038/nature09164

Bose N, Ogawa A, von Reuss SH, et al (2012) Complex Small-Molecule Architectures Regulate Phenotypic Plasticity in a Nematode. Angew Chem-ger Edit 124:12606–12611. 10.1002/ange.201206797

Brodie ED, Brodie ED (1990) Tetrodotoxin Resistance in Garter Snakes: An Evolutionary Response of Predators to Dangerous Prey. Evolution 44:651–659. 10.1111/j.1558-5646.1990.tb05945.x

Brodie ED, Feldman CR, Hanifin CT, et al (2005) Parallel Arms Races between Garter Snakes and Newts Involving Tetrodotoxin as the Phenotypic Interface of Coevolution. J Chem Ecol 31:343–356. 10.1007/s10886-005-1345-x

Chauhan VM, Scurr DJ, Christie T, et al (2017) The physicochemical fingerprint of Necator americanus. Plos Neglect Trop D 11:e0005971. 10.1371/journal.pntd.0005971

Chen J, Caswell-Chen EP (2004) Facultative Vivipary is a Life-History Trait in Caenorhabditis elegans. J Nematol 36:107–13

Ciche TA, Kim K, Kaufmann-Daszczuk B, et al (2008) Cell Invasion and Matricide during Photorhabdus luminescens Transmission by Heterorhabditis bacteriophora Nematodes. Appl Environ Microbiol 74:2275–2287. 10.1128/aem.02646-07

Corcoran AJ, Barber JR, Conner WE (2009) Tiger moth jams bat sonar. Sci (N York, NY) 325:325–7. 10.1126/science.1174096

Crean AJ, Marshall DJ (2009) Coping with environmental uncertainty: dynamic bet hedging as a maternal effect. Philos Trans R Soc B: Biol Sci 364:1087–1096. 10.1098/rstb.2008.0237

Danecek P, Bonfield JK, Liddle J, et al (2021) Twelve years of SAMtools and BCFtools. GigaScience 10:giab008. 10.1093/gigascience/giab008

Demir İ, Demirbağ Z, Erbaş Z (2017) Isolation and Characterization of a Parasitic Nematode, Oscheius myriophila (Nematoda: Rhabditida), Associated with European Mole Cricket, Gryllotalpa gryllotalpa (Orthoptera: Gryllotalpidae). Hacet J Biol Chem 2:197–203. 10.15671/hjbc.2017.152

Dockendorff TC, Estrem B, Reed J, et al (2022) The nematode Oscheius tipulae as a genetic model for programmed DNA elimination. Curr Biol 32:5083–5098.e6. 10.1016/j.cub.2022.10.043

Edgar RC (2004) MUSCLE: multiple sequence alignment with high accuracy and high throughput. Nucleic Acids Res 32:1792–1797. 10.1093/nar/gkh340

Eren GG, Böger L, Roca M, et al (2026) Predatory aggression evolved through adaptations to noradrenergic circuits. Nature 1–10. 10.1038/s41586-025-10009-x

Ettema CH (1998) Soil nematode diversity: species coexistence and ecosystem function. J Nematol 30:159–69

Feldman CR, Brodie ED, Brodie ED, Pfrender ME (2009) The evolutionary origins of beneficial alleles during the repeated adaptation of garter snakes to deadly prey. Proc Natl Acad Sci 106:13415–13420. 10.1073/pnas.0901224106

Felix M-A (2006) Oscheius tipulae. WormBook 1–8. 10.1895/wormbook.1.119.1

Félix M-A, Ailion M, Hsu J-C, et al (2018) Pristionchus nematodes occur frequently in diverse rotting vegetal substrates and are not exclusively necromenic, while Panagrellus redivivoides is found specifically in rotting fruits. Plos One 13:e0200851. 10.1371/journal.pone.0200851

Félix M-A, Duveau F (2012) Population dynamics and habitat sharing of natural populations of Caenorhabditis elegans and C. briggsae. Bmc Biol 10:59. 10.1186/1741-7007-10-59

Ghavamabad RG, Talebi AA, Mehrabadi M, et al (2021) First record of Oscheius myriophilus (Poinar, 1986) (Rhabditida: Rhabditidae) from Iran; and its efficacy against two economic forest trees pests, Cydalima perspectalis (Walker, 1859) (Lepidoptera: Crambidae) and Hyphantria cunea (Drury, 1773) (Lepidoptera: Erebidae) in laboratory condition. J Nematol 53:1–16. 10.21307/jofnem-2021-035

Giblin-Davis RM, Kanzaki N, Davies KA (2013) Nematodes that Ride Insects: Unforeseen Consequences of Arriving Species. Fla Èntomol 96:770–780. 10.1653/024.096.0310

Harry CJ, Messar SM, Ragsdale EJ (2022) Comparative reconstruction of the predatory feeding structures of the polyphenic nematode Pristionchus pacificus. Evol Dev 24:16–36. 10.1111/ede.12397

Herrmann M, Kienle S, Rochat J, et al (2010) Haplotype diversity of the nematode Pristionchus pacificus on Réunion in the Indian Ocean suggests multiple independent invasions. Biol J Linn Soc 100:170–179. 10.1111/j.1095-8312.2010.01410.x

Herrmann M, Mayer WE, Hong RL, et al (2007) The Nematode Pristionchus pacificus (Nematoda: Diplogastridae) Is Associated with the Oriental Beetle Exomala orientalis (Coleoptera: Scarabaeidae) in Japan. Zool Sci 24:883–889. 10.2108/zsj.24.883

Hiramatsu F, Goetting DL, Kotowska AM, et al (2026) Contact-based kin discrimination is associated with specific surface lipids in the cannibalistic nematode Pristionchus pacificus. 10.64898/2026.01.13.699210

Hiramatsu F, Lightfoot JW (2023) Kin-recognition and predation shape collective behaviors in the cannibalistic nematode Pristionchus pacificus. PLOS Genet 19:e1011056. 10.1371/journal.pgen.1011056

Hong RL, Sommer RJ (2006) Chemoattraction in Pristionchus Nematodes and Implications for Insect Recognition. Curr Biol 16:2359–2365. 10.1016/j.cub.2006.10.031

Hong RL, Witte H, Sommer RJ (2008) Natural variation in Pristionchus pacificus insect pheromone attraction involves the protein kinase EGL-4. Proc National Acad Sci 105:7779–7784. 10.1073/pnas.0708406105

Hoogen J van den, Geisen S, Routh D, et al (2019) Soil nematode abundance and functional group composition at a global scale. Nature 572:194–198. 10.1038/s41586-019-1418-6

Ichiishi K, Ekino T, Kanzaki N, Shinya R (2021) Thick cuticles as an anti-predator defence in nematodes. Nematology 24:11–20. 10.1163/15685411-bja10107

Ishita Y, Chihara T, Okumura M (2021) Different combinations of serotonin receptors regulate predatory and bacterial feeding behaviors in the nematode Pristionchus pacificus. G3 Genes Genomes Genetics 11:jkab011-. 10.1093/g3journal/jkab011

Kanzaki N, Giblin-Davis RM (2025) Practical Plant Nematology. 125–163. 10.1079/9781836990413.0007

Kanzaki N, Herrmann M, Weiler C, et al (2021) Nine new Pristionchus (Nematoda: Diplogastridae) species from China. Zootaxa 4943:1–66. 10.11646/zootaxa.4943.1.1

Koneru SL, Salinas H, Flores GE, Hong RL (2016) The bacterial community of entomophilic nematodes and host beetles. Mol Ecol 25:2312–2324. 10.1111/mec.13614

Kotowska AM, Hiramatsu F, Alexander MR, et al (2025) Surface Lipids in Nematodes are Influenced by Development and Species-specific Adaptations. J Am Chem Soc 147:6439–6449. 10.1021/jacs.4c12519

Lenuzzi M, Witte H, Riebesell M, et al (2023) Influence of environmental temperature on mouth-form plasticity in Pristionchus pacificus acts through daf-11-dependent cGMP signaling. J Exp Zoöl Part B: Mol Dev Evol 340:214–224. 10.1002/jez.b.23094

Levis NA, Ragsdale EJ (2025) Genomic parallelism defines repeated evolution of an inducible offense. Sci Adv 11:eadw9964. 10.1126/sciadv.adw9964

Li H, Durbin R (2010) Fast and accurate long-read alignment with Burrows–Wheeler transform. Bioinformatics 26:589–595. 10.1093/bioinformatics/btp698

Lightfoot JW, Dardiry M, Kalirad A, et al (2021) Sex or cannibalism: Polyphenism and kin recognition control social action strategies in nematodes. Sci Adv 7:eabg8042. 10.1126/sciadv.abg8042

Lightfoot JW, Wilecki M, Okumura M, Sommer RJ (2016) Assaying Predatory Feeding Behaviors in *Pristionchus* and Other Nematodes. J Vis Exp. 10.3791/54404

Lightfoot JW, Wilecki M, Rödelsperger C, et al (2019) Small peptide–mediated self-recognition prevents cannibalism in predatory nematodes. Science 364:86–89. 10.1126/science.aav9856

Mayer WE, Herrmann M, Sommer RJ (2007) Phylogeny of the nematode genus Pristionchus and implications for biodiversity, biogeography and the evolution of hermaphroditism. BMC Evol Biol 7:104. 10.1186/1471-2148-7-104

McGaughran A, Rödelsperger C, Grimm DG, et al (2016) Genomic Profiles of Diversification and Genotype–Phenotype Association in Island Nematode Lineages. Mol Biol Evol 33:2257–2272. 10.1093/molbev/msw093

Meyer JM, Baskaran P, Quast C, et al (2017) Succession and dynamics of Pristionchus nematodes and their microbiome during decomposition of Oryctes borbonicus on La Réunion Island. Environ Microbiol 19:1476–1489. 10.1111/1462-2920.13697

Moreno E, Lightfoot JW, Lenuzzi M, Sommer RJ (2019) Cilia drive developmental plasticity and are essential for efficient prey detection in predatory nematodes. Proc Royal Soc B 286:20191089. 10.1098/rspb.2019.1089

Mosser T, Matic I, Leroy M (2011) Bacterium-Induced Internal Egg Hatching Frequency Is Predictive of Life Span in Caenorhabditis elegans Populations. Appl Environ Microbiol 77:8189–8192. 10.1128/aem.06357-11

O’Connell KF (2022) Cryopreservation of C. elegans and Other Nematodes with Dimethyl Sulfoxide and Trehalose. Methods Mol Biol (Clifton, NJ) 2468:43–49. 10.1007/978-1-0716-2181-3_3

Okumura M, Wilecki M, Sommer RJ (2017) Serotonin Drives Predatory Feeding Behavior via Synchronous Feeding Rhythms in the Nematode Pristionchus pacificus. G3 Genes Genomes Genetics 7:3745–3755. 10.1534/g3.117.300263

Philippi T, Seger J (1989) Hedging one’s evolutionary bets, revisited. Trends Ecol Evol 4:41–44. 10.1016/0169-5347(89)90138-9

Pickett CL, Kornfeld K (2013) Age-related degeneration of the egg-laying system promotes matricidal hatching in Caenorhabditis elegans. Aging Cell 12:544–553. 10.1111/acel.12079

Piskobulu V, Athanasouli M, Witte H, et al (2025) High Nutritional Conditions Influence Feeding Plasticity in Pristionchus pacificus and Render Worms Non-Predatory. J Exp Zoöl Part B, Mol Dev Evol 344:94–111. 10.1002/jez.b.23284

Poinar Jr (1986) Rhabditis myriophila sp.n. (Rhabditidae: Rhabditida), associated with the millipede, Oxidis gracilis (Polydesmida: Diplopoda). Proceedings of the Helminthological Society of Washington, Vol. 53,:232–236 ref. 5

Polis GA (1981) The Evolution and Dynamics of Intraspecific Predation. Annu Rev Ecol Syst 12:225–251. 10.1146/annurev.es.12.110181.001301

Polis GA, Holt RD (1992) Intraguild predation: The dynamics of complex trophic interactions. Trends Ecol Evol 7:151–154. 10.1016/0169-5347(92)90208-s

Polis GA, Myers CA, Holt RD (1989) The Ecology and Evolution of Intraguild Predation: Potential Competitors That Eat Each Other. Annu Rev Ecol Syst 20:297–330. 10.1146/annurev.es.20.110189.001501

Qing X, Zhang YM, Sun S, et al (2024) Phylogenomic Insights into the Evolution and Origin of Nematoda. Syst Biol syae073. 10.1093/sysbio/syae073

Quach KT, Chalasani SH (2022) Flexible reprogramming of Pristionchus pacificus motivation for attacking Caenorhabditis elegans in predator-prey competition. Curr Biol. 10.1016/j.cub.2022.02.033

Ragsdale EJ, Müller MR, Rödelsperger C, Sommer RJ (2013) A Developmental Switch Coupled to the Evolution of Plasticity Acts through a Sulfatase. Cell 155:922–933. 10.1016/j.cell.2013.09.054

Renahan T, Lo W, Werner MS, et al (2021) Nematode biphasic ‘boom and bust’ dynamics are dependent on host bacterial load while linking dauer and mouth-form polyphenisms. Environ Microbiol. 10.1111/1462-2920.15438

Roca M, Eren GG, Böger L, et al (2026) Evolution of sensory systems underlies the emergence of predatory feeding behaviors in nematodes. Proc Natl Acad Sci United States Am 123:e2514172123. 10.1073/pnas.2514172123

Rödelsperger C, Meyer JM, Prabh N, et al (2017) Single-Molecule Sequencing Reveals the Chromosome-Scale Genomic Architecture of the Nematode Model Organism Pristionchus pacificus. Cell Reports 21:834–844. 10.1016/j.celrep.2017.09.077

Rödelsperger C, Neher RA, Weller AM, et al (2014) Characterization of Genetic Diversity in the Nematode Pristionchus pacificus from Population-Scale Resequencing Data. Genetics 196:1153–1165. 10.1534/genetics.113.159855

Rödelsperger C, Röseler W, Athanasouli M, et al (2024) Genome Assembly of the Nematode Rhabditoides Inermis From a Complex Microbial Community. Genome Biol Evol 16:evae230. 10.1093/gbe/evae230

Rödelsperger C, Röseler W, Prabh N, et al (2018) Phylotranscriptomics of Pristionchus Nematodes Reveals Parallel Gene Loss in Six Hermaphroditic Lineages. Curr Biol 28:3123–3127.e5. 10.1016/j.cub.2018.07.041

Rosa PMG de la, Thomson M, Trivedi U, et al (2020) A telomere-to-telomere assembly of Oscheius tipulae and the evolution of rhabditid nematode chromosomes. G3: GenesGenomesGenet 11:jkaa020. 10.1093/g3journal/jkaa020

Ryan WBF, Carbotte SM, Coplan JO, et al (2009) Global Multi-Resolution Topography synthesis: GLOBAL MULTI-RESOLUTION TOPOGRAPHY SYNTHESIS. Geochem, Geophys, Geosystems 10:n/a-n/a. 10.1029/2008gc002332

Schliep KP (2010) phangorn: phylogenetic analysis in R. Bioinformatics 27:592–593. 10.1093/bioinformatics/btq706

Schmitz O (2017) Predator and prey functional traits: understanding the adaptive machinery driving predator–prey interactions. F1000Research 6:1767. 10.12688/f1000research.11813.1

Sommer Ralf J, Carta LK, Kim S-Y, Sternberg PW (1996) Morphological, genetic and molecular description of Pristionchus pacificus sp. n.(Nematoda: Neodiplogastridae). Fundam appl Nematol (6):511–521

Sommer RJ (2025) Pristionchus – Beetle associations: Towards a new natural history. J Invertebr Pathol 209:108243. 10.1016/j.jip.2024.108243

Sommer RJ, Lightfoot JW (2022) Nematodes as Model Organisms. 1–23. 10.1079/9781789248814.0001

Sudhaus W, Hooper DJ (1994) Rhabditis (Oscheius) Guentheri1 Sp.N., an Unusual Species With Reduced Posterior Ovary, With Observations On the Dolichura and Insectivora Groups (Nematoda: Rhabditidae). Nematologica 40:508–533. 10.1163/003525994x00391

Werner MS, Claaßen MH, Renahan T, et al (2018) Adult Influence on Juvenile Phenotypes by Stage-Specific Pheromone Production. Iscience 10:123–134. 10.1016/j.isci.2018.11.027

Werner MS, Sieriebriennikov B, Loschko T, et al (2017) Environmental influence on Pristionchus pacificus mouth form through different culture methods. Sci Rep-uk 7:7207. 10.1038/s41598-017-07455-7

Wilecki M, Lightfoot JW, Susoy V, Sommer RJ (2015) Predatory feeding behaviour in Pristionchus nematodes is dependent on phenotypic plasticity and induced by serotonin. J Exp Biol 218:1306–1313. 10.1242/jeb.118620

Yeates GW, Bongers T, Goede RGD, et al (1993) Feeding habits in soil nematode families and genera-an outline for soil ecologists. J Nematol 25:315–31

