## Supplemental Files for "Intraguild predation and life-history variation in a natural nematode predator-prey system"

**Figure S1**

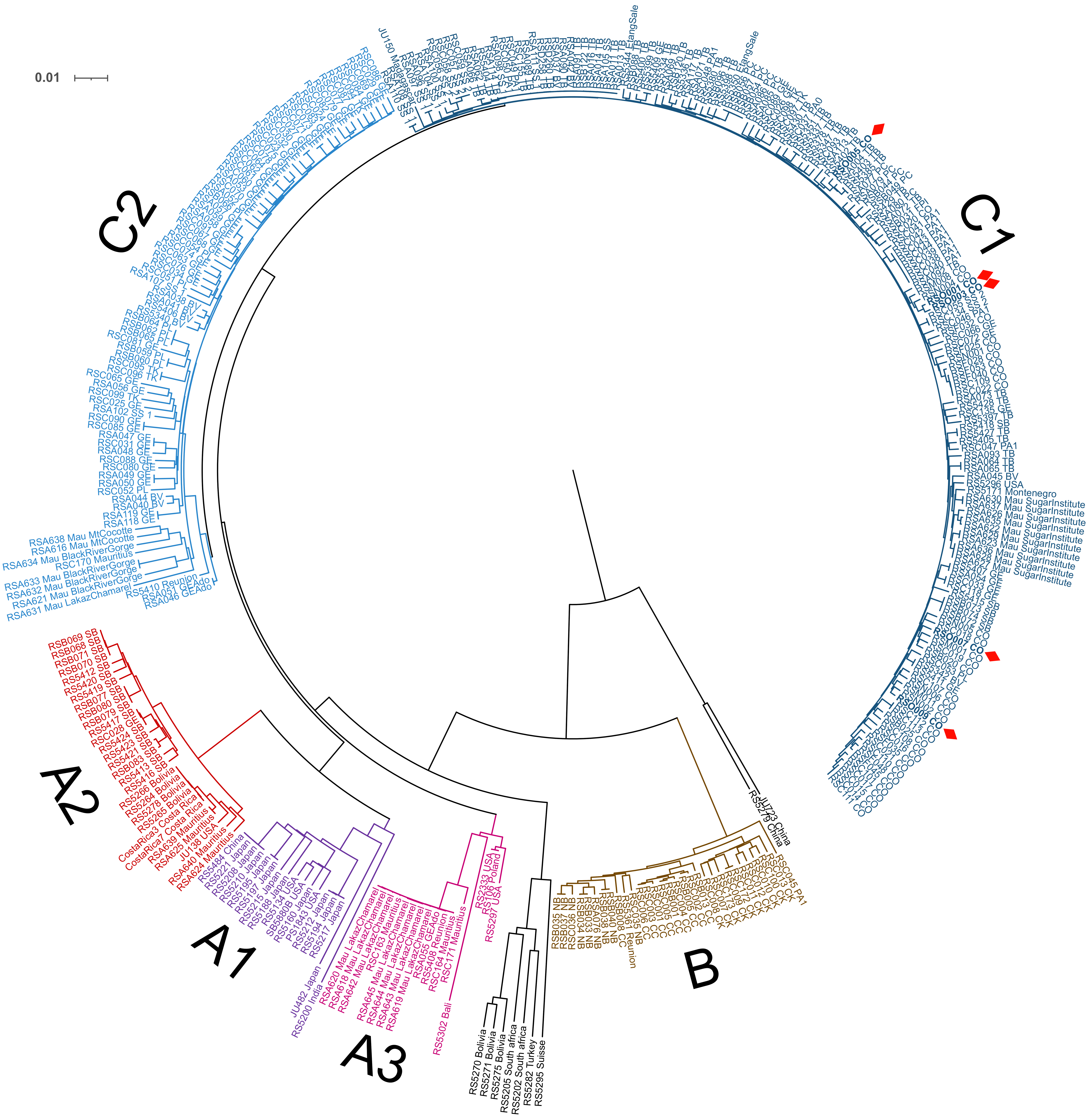

**Figure S2**

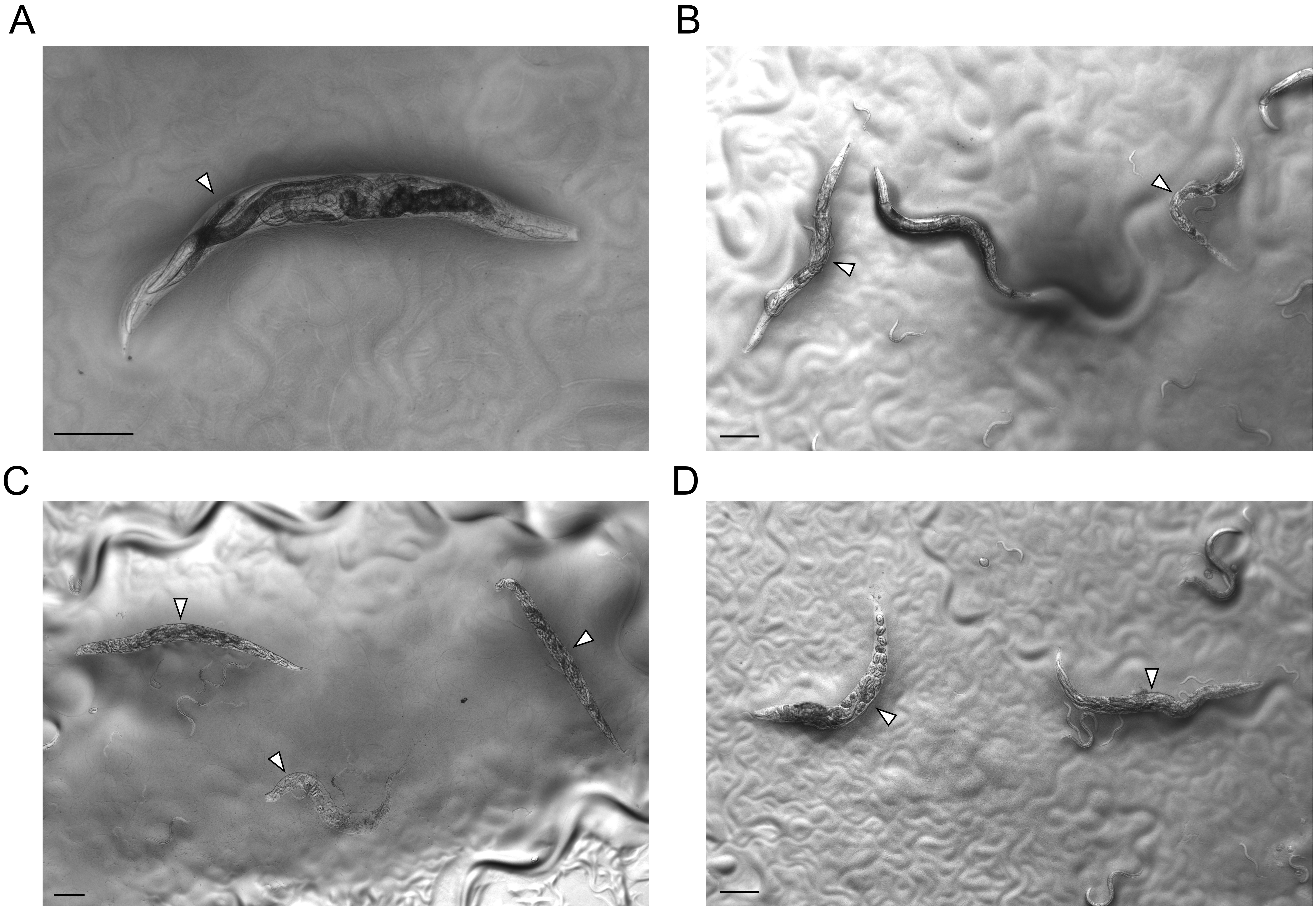

### Figure S3

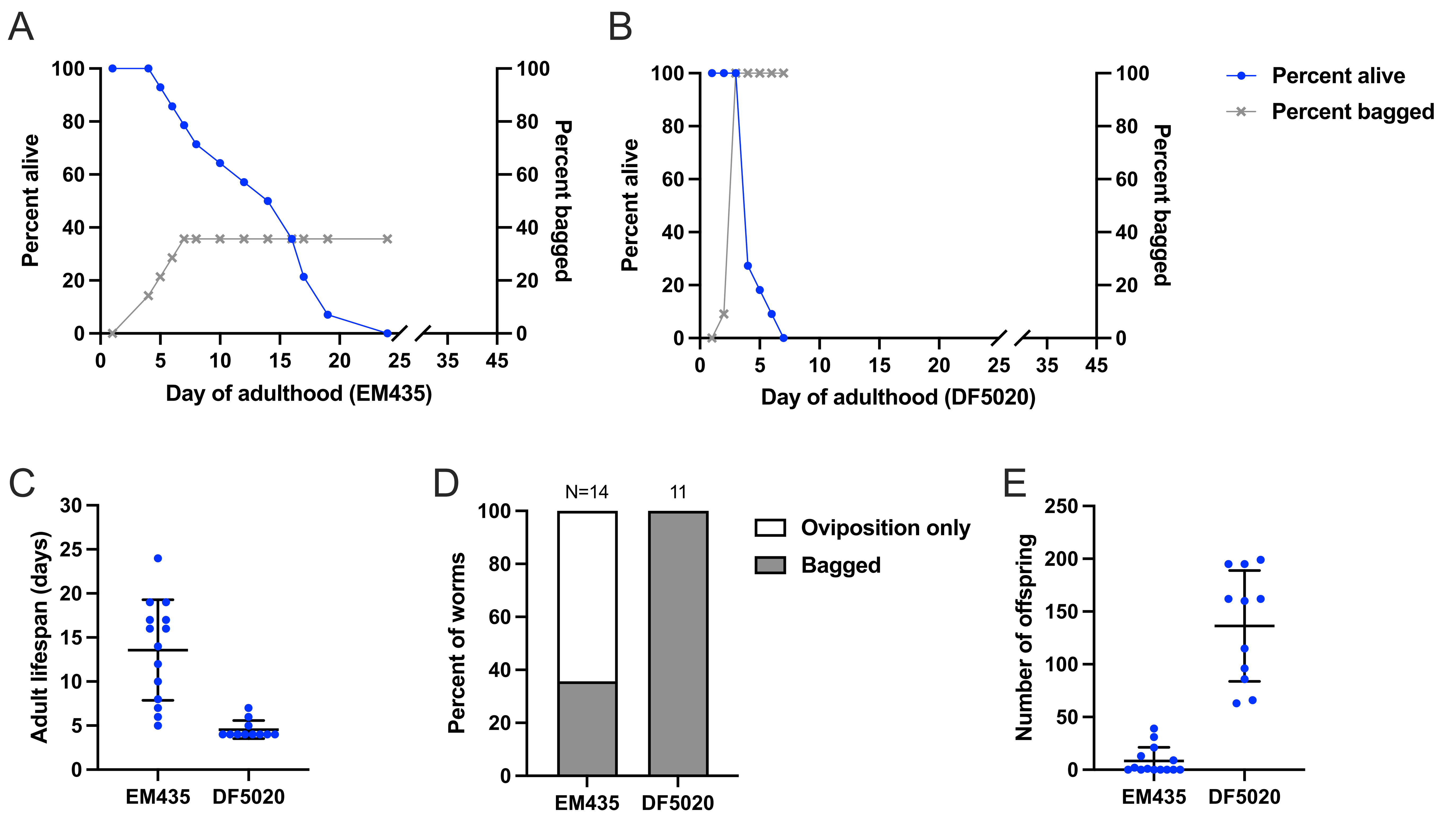
A
Percent alive
Percent bagged
Day of adulthood (EM435)
B
Percent alive
Percent bagged
Day of adulthood (DF5020)
Percent alive
Percent bagged
C
Adult lifespan (days)
EM435
DF5020
D
N=14
11
Percent of worms
Oviposition only
Bagged
EM435
DF5020
E
Number of offspring
EM435
DF5020
